# Ensembler: Enabling high-throughput molecular simulations at the superfamily scale

**DOI:** 10.1101/018036

**Authors:** Daniel L. Parton, Patrick B. Grinaway, Sonya M. Hanson, Kyle A. Beauchamp, John D. Chodera

## Abstract

The rapidly expanding body of available genomic and protein structural data provides a rich resource for understanding protein dynamics with biomolecular simulation. While computational infrastructure has grown rapidly, simulations on an *omics* scale are not yet widespread, primarily because software infrastructure to enable simulations at this scale has not kept pace. It should now be possible to study protein dynamics across entire (super)families, exploiting both available structural biology data and conformational similarities across homologous proteins. Here, we present a new tool for enabling high-throughput simulation in the genomics era. **Ensembler** takes any set of sequences—from a single sequence to an entire superfamily— and shepherds them through various stages of modeling and refinement to produce simulation-ready structures. This includes comparative modeling to all relevant PDB structures (which may span multiple conformational states of interest), reconstruction of missing loops, addition of missing atoms, culling of nearly identical structures, assignment of appropriate protonation states, solvation in explicit solvent, and refinement and filtering with molecular simulation to ensure stable simulation. The output of this pipeline is an ensemble of structures ready for subsequent molecular simulations using computer clusters, supercomputers, or distributed computing projects like Folding@home. **Ensembler** thus automates much of the time consuming process of preparing protein models suitable for simulation, while allowing scalability up to entire superfamilies. A particular advantage of this approach can be found in the construction of kinetic models of conformational dynamics—such as Markov state models (MSMs)—which benefit from a diverse array of initial configurations that span the accessible conformational states to aid sampling. We demonstrate the power of this approach by constructing models for all catalytic domains in the human tyrosine kinase family, using all available kinase catalytic domain structures from any organism as structural templates.

**Ensembler** is free and open source software licensed under the GNU General Public License (GPL) v2. It is compatible with Linux and OS X. The latest release can be installed via the conda package manager, and the latest source can be downloaded from https://github.com/choderalab/ensembler.

## I. INTRODUCTION

Recent advances in genomics and structural biology have helped generate an enormous wealth of protein data at the level of amino-acid sequence and three-dimensional structure. However, proteins typically exist as an ensemble of thermally accessible conformational states, and static structures provide only a snapshot of their rich dynamical behavior. Many functional properties—such as the ability to bind small molecules or interact with signaling partners—require transitions between states, encompassing anything from reorganization of sidechains at binding interfaces to domain motions to large scale folding-unfolding events. Drug discovery could also benefit from a more extensive consideration of protein dynamics, whereby small molecules might be selected based on their predicted ability to bind and trap a protein target in an inactive state [1].

Molecular dynamics (MD) simulations have the capability, in principle, to describe the time evolution of a protein in atomistic detail, and have proven themselves to be a useful tool in the study of protein dynamics. A number of mature software packages and force fields are now available, and much recent progress has been driven by advances in computing architecture. For example, many MD packages are now able to exploit GPUs [2, 3], which provide greatly improved simulation efficiency per unit cost relative to CPUs, while distributed computing platforms such as Folding@home [4], Copernicus [5, 6], and GPUGrid [7], allow scalability on an unprecedented level. In parallel, methods for building human-understandable models of protein dynamics from noisy simulation data, such as Markov state modeling (MSM) approaches, are now reaching maturity [8–10]. MSM methods in particular have the advantage of being able to aggregate data from multiple independent MD trajectories, facilitating parallelization of production simulations and thus greatly alleviating overall computational cost. There also exist a number of mature software packages for comparative modeling of protein structures, in which a target protein sequence is modeled using one or more structures as templates [11, 12]. One such piece of software, MODELLER, has also been used recently to study protein allostery by generating and refining configurational models, sampled by interpolating between two user-defined metastable structures [13].

However, it remains difficult for researchers to exploit the full variety of available protein sequence and structural data in simulation studies, largely due to limitations in software architecture. For example, the set up of a biomolecular simulation is typically performed manually, encompassing a series of fairly standard (yet time-consuming) steps such as the choice of protein sequence construct and starting structure(s), addition of missing residues and atoms, solvation with explicit water and counterions (and potentially buffer components and cosolvents), choice of simulation parameters (or parameterization schemes for components where parameters do not yet exist), system relaxation with energy minimization, and one or more short preparatory MD simulations to equilibrate the system and relax the simulation cell. Due to the laborious and manual nature of this process, simulation studies typically consider only one or a few proteins and starting configurations. Worse still, studies (or collections of studies) that do consider multiple proteins often suffer from the lack of consistent best practices in this preparation process, making comparisons between related proteins unnecessarily difficult.

The ability to fully exploit the large quantity of available protein sequence and structural data in biomolecular simulation studies could open up many interesting avenues for research, enabling the study of entire protein families or superfamilies within a single organism or across multiple organisms. The similarity between members of a given protein family could be exploited to generate arrays of conformational models, which could be used as starting configurations to aid sampling in MD simulations. This approach would be highly beneficial for many MD methods, such as MSM construction, which require global coverage of the conformational landscape to realize their full potential, and would also be particularly useful in cases where structural data is present for only a subset of the members of a protein family. It would also aid in studying protein families known to have multiple metastable conformations—such as kinases—for which the combined body of structural data for the family may cover a large range of these conformations, while the available structures for any individual member might encompass only one or two distinct conformations.

Here, we present the first steps toward bridging the gap between biomolecular simulation software and *omics-* scale sequence and structural data: a fully automated open source framework for building simulation-ready protein models in multiple conformational substates scalable from single sequences to entire superfamilies. **Ensembler** provides functions for selecting target sequences and homologous template structures, and (by interfacing with a number of external packages) performs pairwise alignments, comparative modeling of target-template pairs, and several stages of model refinement. As an example application, we have constructed models for the entire set of human tyrosine kinase (TK) catalytic domains, using all available structures of protein kinase domains (from any species) as templates. This results in a total of almost 400,000 models, and we demonstrate that these provide wide-ranging coverage of known functionally relevant conformations. By using these models as starting configurations for highly parallel MD simulations, we expect their structural diversity to greatly aid in sampling of conformational space. We further suggest that models with high target-template sequence identity are the most likely to represent native metastable states, while lower sequence identity models would aid in sampling of more distant regions of accessible phase space. It is also important to note that some models (especially low sequence identity models) may not represent natively accessible conformations. However, MSM methods benefit from the ability to remove outlier MD trajectories which start from non-natively accessible conformations, and which would thus be unconnected with the phase space sampled in other trajectories. These methods essentially identify the largest subset of Markov nodes which constitute an ergodic network [14, 15].

We anticipate that **Ensembler** will prove to be useful in a number of other ways. For example, the generated models could represent valuable data sets even without subsequent production simulation, allowing exploration of the conformational diversity present within the available structural data for a given protein family. Furthermore, the automation of simulation set up provides an excellent opportunity to make concrete certain “best practices”, such as the choice of simulation parameters.

## II. DESIGN AND IMPLEMENTATION

**Ensembler** is written in Python, and can be used via a command-line tool (ensembler) or via a flexible Python API to allow integration of its components into other applications. All command-line and API information in this article refers to the version 1.0.2 release of Ensem https://github.com/choderalab/ensembler/tree/v1.0.2”bler. Up-to-date documentation can be found at ensembler.readthedocs.org.

The **Ensembler** modeling pipeline comprises a series of stages which are performed in a defined order. A visual overview of the pipeline is shown in Fig. 1. The various stages of this pipeline are described in detail below.

**FIG. 1.**
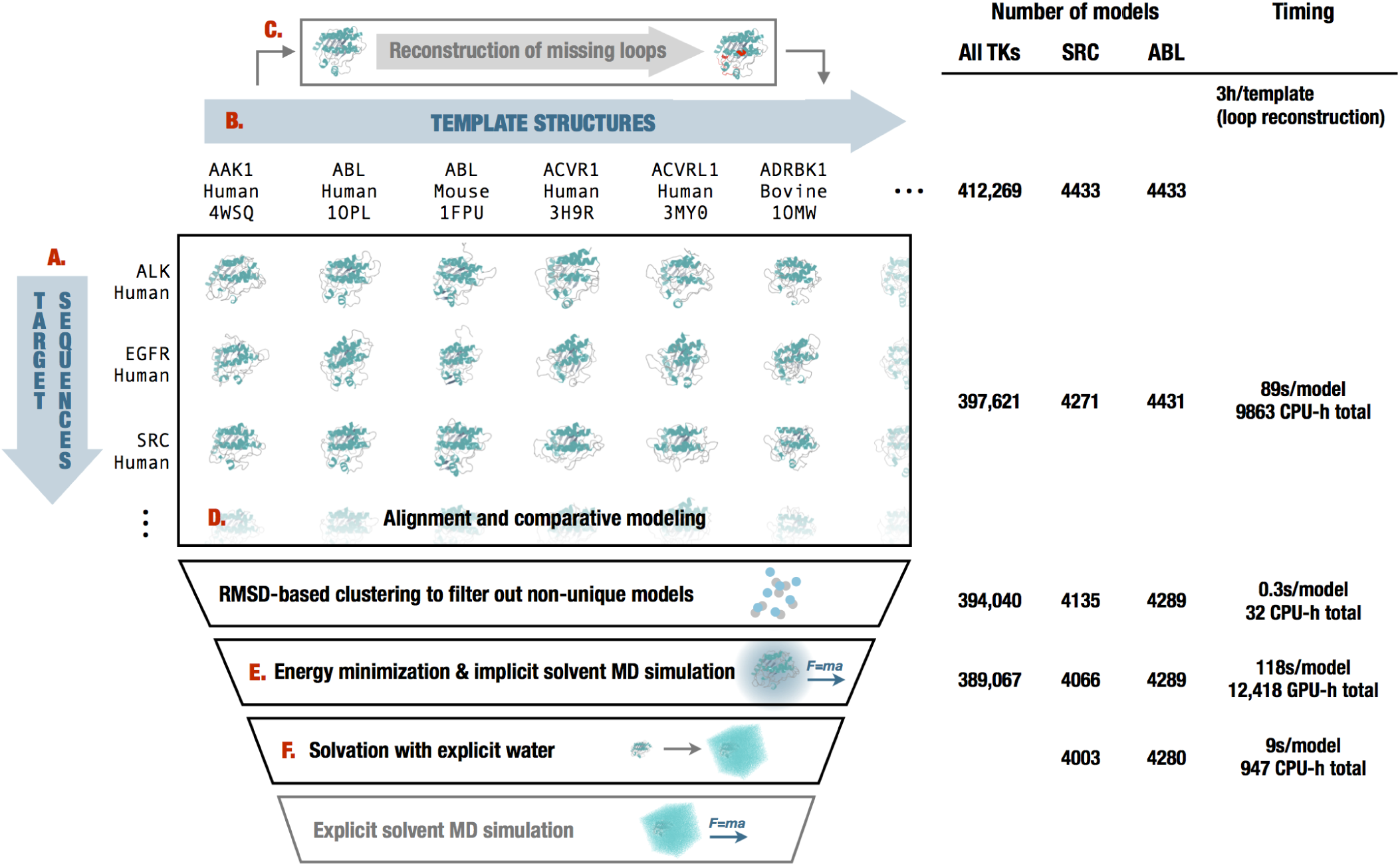
Diagrammatic representation of the stages of the Ensembler pipeline and illustrative statistics for modeling all human tyrosine kinase catalytic domains. On the left, the various stages of the **Ensembler** pipeline are shown. The red labels indicate the corresponding text description provided for each stage in the Design and Implementation section. On the right, the number of viable models surviving each stage of the pipeline is shown for the 93 target TK domains and for two representative individual TK domains *(SRC* and *ABL).* Typical timings on a computer cluster (containing Intel Xeon E5-2665 2.4GHz hyperthreaded processors and NVIDIA GTX-680 or GTX-Titan GPUs) is reported to illustrate resource requirements per model for modeling the entire set of tyrosine kinases. Note that *CPU-h* denotes the number of hours consumed by the equivalent of a single CPU hyperthread and *GPU-h* on a single GPU—parallel execution via MPI reduces wall clock time nearly linearly.

### A. Target selection and retrieval

The first stage entails the selection of a set of *target* protein sequences—the sequences for which the user is interested in generating simulation-ready structural models. This may be a single sequence—such as a full-length protein or a construct representing a single domain—or a collection of sequences, such as a particular domain from an entire family of proteins. The output of this stage is a FASTA-formatted text file containing the desired target sequences with corresponding arbitrary identifiers.

The `ensembler` command-line tool allows targets to be selected from UniProt—a freely accessible resource for protein sequence and functional data (uniprot.org) [16]— via a UniProt search query. To retrieve target sequences from UniProt, the subcommand `gather_targets` is used with the `--query flag` followed by a UniProt query string conforming to the same syntax as the search function available on the UniProt website. For example, `--query ‘mnemonic:SRC_HUMAN’` would select the full-length human Src sequence, while the query shown in Box 1 would select all human tyrosine protein kinases which have been reviewed by a human curator. In this way, the user may select a single protein, many proteins, or an entire superfamily from UniProt. The program outputs a FASTAfile, setting the UniProt mnemonic (e.g. SRC_HUMAN) as the identifier for each target protein.

#### Box 1.

##### Ensembler command-line functions used to select targets and templates.

The commands retrieve target and template data by querying UniProt. The query string provided to the `gather_targets` command selects all human tyrosine protein kinases which have been reviewed by a curator, while the query string provided to the gather_templates command selects all reviewed protein kinases of any species. The `--uniprot_domain_regex flag` is used to select a subset of the domains belonging to the returned UniProt protein entries, by matching the domain annotations against a given regular expression. In this example, domains of type “Protein kinase”, “Protein kinase 1”, and “Protein kinase 2” were selected, while excluding many other domain types such as “Protein kinase; truncated”, “Protein kinase; inactive”, “SH2”, “SH3”, etc. Target selection simply entails the selection of sequences corresponding to each matching UniProt domain. Template selection entails the selection of the sequences and structures of any PDB entries corresponding to the matching UniProt domains.

~~~
ensembler gather_targets --query ‘family:”tyr protein kinase family” AND organism:”homo sapiens” AND reviewed:yes’
                        --uniprot_domain_regex ‘^Protein kinase(?!; truncated)(?!; inactive)’
ensembler gather_templates --gather_from uniprot --query ‘domain:”Protein kinase” AND reviewed:yes’
                          --uniprot_domain_regex ‘^Protein kinase(?!; truncated)(?!; inactive)’
~~~

In many cases, it will be desirable to build models of an isolated protein domain, rather than the full-length protein. The `gather_targets` subcommand allows protein domains to be selected from UniProt data by passing a regular expression string to the `--uniprot_domain_regex` flag. For example, the above `--query flag` for selecting all human protein kinases returns UniProt entries with domain annotations including “Protein kinase”, “Protein kinase 1”, “Protein kinase 2”, “Protein kinase; truncated”, “Protein kinase; inactive”, “SH2”, “SH3”, etc. The regular expression shown in Box 1 selects only domains of the first three types. If the `--uniprot_domain_regex` flag is used, target identifiers are set with the form `[UniProt mnemonic]_D[domain index]`, where the latter part represents a 0-based index for the domain—necessary because a single target protein may contain multiple domains of interest (e.g. JAK1_HUMAN_D0, JAK1_HUMAN_D1).

Target sequences can also be defined manually (or from another program) by providing a FASTA-formatted text file containing the desired target sequences with corresponding arbitrary identifiers.

### B. Template selection and retrieval

**Ensembler** uses comparative modeling to build models, and as such requires a set of structures to be used as templates. The second stage thus entails the selection of templates and storage of associated sequences, structures, and identifiers. These templates can be specified manually, or using the ensembler gather_templates subcommand to automatically select templates based on a search of the Protein Data Bank (PDB) or UniProt. A recommended approach is to select templates from UniProt which belong to the same protein family as the targets, guaranteeing some degree of homology between targets and templates.

The ensembler gather_templates subcommand provides methods for selecting template structures from either UniProt or the PDB (http://www.rcsb.org/pdb), specified by the `--gather_from` flag. Both methods select templates at the level of PDB chains—a PDB structure containing multiple chains with identical sequence spans (e.g. for crystal unit cells with multiple asymmetric units) would thus give rise to multiple template structures.

Selection of templates from the PDB simply requires passing a list of PDB IDs as a comma-separated string, e.g. `--query 2H8H,1Y57.` Specific PDB chain IDs can optionally also be selected via the --chainids flag. The program retrieves structures from the PDB server, as well as associated data from the SIFTS service (www.ebi.ac.uk/pdbe/docs/sifts) [17], which provides residue-level mappings between PDB and UniProt entries. The SIFTS data is used to extract template sequences, retaining only residues which are resolved and match the equivalent residue in the UniProt sequence—non-wildtype residues are thus removed from the template structures. Furthermore, PDB chains with less than a given percentage of resolved residues (default: 70%) are filtered out. Sequences are stored in a FASTA file, with identifiers of the form `[UniProt mnemonic]_D[UniProt domain index]_[PDB ID]_[PDB chain ID]`, e.g. `SRC_HUMAN_D0_2H8H_A`. Matching residues then extracted from the original coordinate files and stored as PDB-format coordinate files.

Selection of templates from UniProt proceeds in a similar fashion as for target selection; the `--query flag` is used to select full-length proteins from UniProt, while the optional `--uniprot_domain_regex` flag allows selection of individual domains with a regular expression string (Box 1). The returned UniProt data for each protein includes a list of associated PDB chains and their residue spans, and this information is used to select template structures, using the same method as for template selection from the PDB. Only structures solved by X-ray crystallography or NMR are selected, thus excluding computer-generated models available from the PDB. If the `--uniprot_domain_regex` flag is used, then templates are truncated at the start and end of the domain sequence.

Templates can also be defined manually. Manual specification of templates simply requires storing the sequences and arbitrary identifiers in a FASTA file, and the structures as PDB-format coordinate files with filenames matching the identifiers in the sequence file. The structure residues must also match those in the sequence file.

### C. Template refinement

Unresolved template residues can optionally be modeled into template structures with the **loopmodel** subcommand, which employs a kinematic closure algorithm provided via the **loopmodel** tool of the Rosetta software suite [18, 19]. We expect that in certain cases, pre-building template loops with Rosetta **loopmodel** prior to the main modeling stage (with MODELLER) may result in improved model quality. Loop remodeling may fail for a small proportion of templates due to spatial constraints imposed by the original structure; the subsequent modeling step thus automatically uses the remodeled version of a template if available, but otherwise falls back to using the non-remodeled version. Furthermore, the Rosetta **loopmodel** program will not model missing residues at the termini of a structure—such residue spans are modeled in the subsequent stage.

### D. Modeling

In the modeling stage, structural models of the target sequence are generated from the template structures, with the goal of modeling the target in a variety of conformations that could be significantly populated under equilibrium conditions.

Modeling is performed using the automodel function of the MODELLER software package [20, 21] to rapidly generate a single model of the target sequence from each template structure. MODELLER uses simulated annealing cycles along with a minimal forcefield and spatial restraints— generally Gaussian interatomic probability densities extracted from the template structure with database-derived statistics determining the distribution width—to rapidly generate candidate structures of the target sequence from the provided template sequence [20, 21].

While MODELLER’s automodel function can generate its own alignments automatically, a standalone function was preferable for reasons of programming convenience. As such, we implemented pairwise alignment functionality using the BioPython `pairwise2` module [22]—which uses a dynamic programming algorithm—with the PAM 250 scoring matrix of Gonnet *et al.* [23]. The alignments are carried out with the **align** subcommand, prior to the modeling step which is carried out with the `build_models` subcommand. The **align** subcommand also writes a list of the sequence identities for each template to a text file, and this can be used to select models from a desired range of sequence identities. The `build_models` subcommand and all subsequent pipeline functions have a **--template_seqid_cutoff** flag which can be used to select only models with sequence identities greater than the given value. We also note that alternative approaches could be used for the alignment stage. For example, multiple sequence alignment algorithms [24], allow alignments to be guided using sequence data from across the entire protein family of interest, while (multiple) structural alignment algorithms such as MODELLER’s **salign** routine [20, 21], PROMALS3D [25], and Expresso and 3DCoffee [26, 27], can additionally exploit structural data. **Ensembler’s** modular architecture facilitates the implementation of alternative alignment approaches, and we plan to implement some of these in future versions, to allow exploration of the influence of different alignment methods on model quality.

Models are output as PDB-format coordinate files. To minimize file storage requirements, **Ensembler** uses the Python **gzip** library to apply compression to all sizeable text files from the modeling stage onwards. The restraints used by MODELLER could potentially be used in alternative additional refinements chemes, and **Ensembler** thus provides a flag (--write_modeller_restraints_file) for optionally saving these restraints to file. This option is turned off by default, as the restraint files are relatively large (e.g. ∼400 kB per model for protein kinase domain targets), and are not expected to be used by the majority of users.

#### Filtering of nearly identical models

Because **Ensembler** treats individual chains from source PDB structures as individual templates, a number of models may be generated with very similar structures if these individual chains are nearly identical in conformation. For this reason, and also to allow users to select for high diversity if they so choose, **Ensembler** provides a way to filter out models that are very similar in RMSD. The cluster subcommand can thus be used to identify models which differ from other models in terms of RMSD distance by a user-specified cutoff. Clustering is performed using the regular spatial clustering algorithm [9], as implemented in the MSM-Builder Python library [14], which uses mdtraj[28] to calculate RMSD (for C_a_ atoms only) with a fast quaternion characteristic polynomial (QCP) [29 -31] implementation. A minimum distance cutoff (which defaults to 0.6 A) is used to retain only a single model per cluster.

### E. Refinement of models

A number of refinement methods have been developed to help guide comparative modeling techniques toward more “native-like” and physically consistent conformations [32, 33], of which MD simulations are an important example. While long-timescale unrestrained MD simulations (on the order of 100 μs) have been found to be ineffective for recapitulating native-like conformations, possibly due to forcefield issues [34], even relatively short simulations can be useful for relaxing structural elements such as sidechain orientation [33].

**Ensembler** thus includes a refinement module, which uses short molecular dynamics simulations to refine the models built in the previous step. As well as improving model quality, this also prepares models for subsequent production MD simulation, including solvation with explicit water molecules, if desired.

Models are first subjected to energy minimization (using the L-BFGS algorithm [35], followed by a short molecular dynamics (MD) simulation with an implicit solvent representation. This is implemented usingthe OpenMM molecular simulation toolkit [2], chosen for its flexible Python API, and high performance GPU-acclerated simulation code. The simulation is run fora default of 100 ps, which in our example applications has been sufficient to filter out poor models (i.e. those with atomic overlaps unresolved by energy minimization, which result in an unstable simulation), as well as helping to relax model conformations. As discussed in the Results section, our example application of the **Ensembler** pipeline to the human tyrosine kinase family indicated that of the models which failed implicit solvent MD refinement, the vast majority failed within the first 1 ps of simulation.

The simulation protocol and default parameter values have been chosen to represent current “best practices” for the refinement simulations carried out here. As such, the simulation is performed using Langevin dynamics, with a default force field choice of Amber99SB-ILDN [36], along with a modified generalized Born solvent model [37] as implemented in the OpenMM package [2]. Any of the other force fields or implicit water models implemented in OpenMM can be specified using the --ff and `--water_model` flags respectively. The simulation length can also be controlled via the `--simlength` flag, and many other important simulation parameters can be controlled from either the API orCLI (via the --api_params flag). The default values are set as follows—timestep: 2 fs; temperature: 300 K; Langevin collision rate: 20 ps^-1^; pH (used by OpenMM for protonation state assignment): 7. We also draw attention to a recent paper which indicates that lower Langevin collision rates may result in faster phase space exploration [38].

### F. Solvation and NPT equilibration

While protein-only models may be sufficient for structural analysis or implicit solvent simulations, **Ensembler** also provides a stage for solvating models with explicit water and performing a round of explicit-solvent MD refinement/equilibration under isothermal-isobaric (NPT) conditions. The solvation step solvates each model for a given target with the same number of waters to facilitate the integration of data from multiple simulations, which is important for methods such as the construction of MSMs. The target number of waters is selected by first solvating each model with a specified padding distance (default: 10 Å), then taking a percentile value from the distribution (default: 68th percentile). This helps to prevent models with particularly long, extended loops—such as those arising from template structures with unresolved termini—from imposing very large box sizes on the entire set of models. The TIP3P water model [39] is used by default, but any of the other explicit water models available in OpenMM, such as TIP4P-Ew [40], can be specified using the `--water_model` flag. Models are resolvated with the target number of waters by first solvating with zero padding, then incrementally increasing the box size and resolvating until the target is exceeded, then finally deleting sufficient waters to match the target value. The explicit solvent MD simulation is also implemented using OpenMM, using the Amber99SB-ILDN force field [36] and TIP3P water [39] by default. The force field, water model, and simulation length can again be specified using the --ff, `--water_model`, and `--simlength` flags respectively. Further simulation parameters can be controlled via the API or via the CLI --api_params flag. Pressure control is performed with a Monte Carlo barostat as implemented in OpenMM, with a default pressure of 1 atm and a period of 50 timesteps. The remaining simulation parameters have default values set to the same as for the implicit solvent MD refinement.

#### Packaging

**Ensembler** provides a packaging module which can be used to prepare models for other uses. The package_models subcommand currently provides functions (specified via the `--package_`for flag) for compressing models in preparation for data transfer, or for organizing them with the appropriate directory and file structure for production simulation on the distributed computing platform Folding@home [4]. The module could easily be extended to add methods for preparing models for other purposes. For example, production simulations could alternatively be run using Copernicus [5, 6]—a framework for performing parallel adaptive MD simulations— or GPUGrid [7]—a distributing computing platform which relies on computational power voluntarily donated by the owners of non dedicated GPU-equipped computers.

#### Other features

##### Tracking provenance information

To aid the user in tracking the provenance of each model, each pipeline function also outputs a metadata file, which helps to link data to the software version used to generate it (both **Ensembler** and its dependencies), and also provides timing and performance information, and other data such as hostname.

##### Rapidly modeling a single template

For users interested in simply using **Ensembler** to rapidly generate a set of models for a single template sequence, **Ensembler** provides a command-line tool quickmodel, which performs the entire pipeline for a single target with a small number of templates. For larger numbers of models (such as entire protein families), modeling time is greatly reduced by using the main modeling pipeline, which is parallelized via MPI, distributing computation across each model (or across each template, in the case of the loop reconstruction code), and scaling (in a “pleasantly parallel” manner) up to the number of models generated.

## III. RESULTS

### Modeling of all human tyrosine kinase catalytic domains

As a first application of **Ensembler**, we have built models for the human TK family. TKs (and protein kinases in general) play important roles in many cellular processes and are involved in a number of types of cancer [41]. For example, a translocation between the TKAbl1 and the pseudokinase Bcr is closely associated with chronic myelogenous leukemia [42], while mutations of Src are associated with colon, breast, prostate, lung, and pancreatic cancers [43]. Protein kinase domains are thought to have multiple accessible metastable conformation states, and much effort is directed at developing kinase inhibitor drugs which bind to and stabilize inactive conformations [44]. Kinases are thus a particularly interesting subject for study with MSM methods [45], and this approach stands to benefit greatly from the ability to exploit the full body of available genomic and structural data within the kinase family, e.g. by generating large numbers of starting configurations to be used in highly parallel MD simulation.

We selected all human TK domains annotated in UniProt as targets, and all available structures of protein kinase domains (of any species) as templates, using the commands shown in Box 1. This returned 93 target sequences and 4433 template structures, giving a total of 412,269 target-template pairs. The templates were derived from 3028 individual PDB entries and encompassed 23 different species, with 3634 template structures from human kinase constructs.

The resultant models are available as part of a supplementary dataset which can be downloaded from the Dryad Digital Repository (DOI: 10.5061/dryad.7fg32).

### Ensembler modeling statistics

Crystallographic structures of kinase catalytic domains generally contain a significant number of missing residues (median 11, mean 14, standard deviation 13, max102) due to the high mobility of several loops (Fig. 2, top), with a number of these missing spans being significant in length (median 5, mean 7, standard deviation 6, max82; Fig.2, bottom). To reduce the reliance on the MODELLER rapid model construction stage to reconstruct very long unresolved loops, unresolved template residues were first remodeled using the loopmodel subcommand. Out of 3666 templates with one or more missing residues, 3134 were successfully remodeled by the Rosetta loop modeling stage (with success defined simply as program termination without error); most remodeling failures were attributable to unsatisfiable spatial constraints imposed by the original template structure. There was some correlation between remodeling failures and the number of missing residues (Fig. 2, top); templates for which remodeling failed had a median of 20 missing residues, compared to a median of 14 missing residues for templates for which remodeling was successful.

**FIG. 2.**
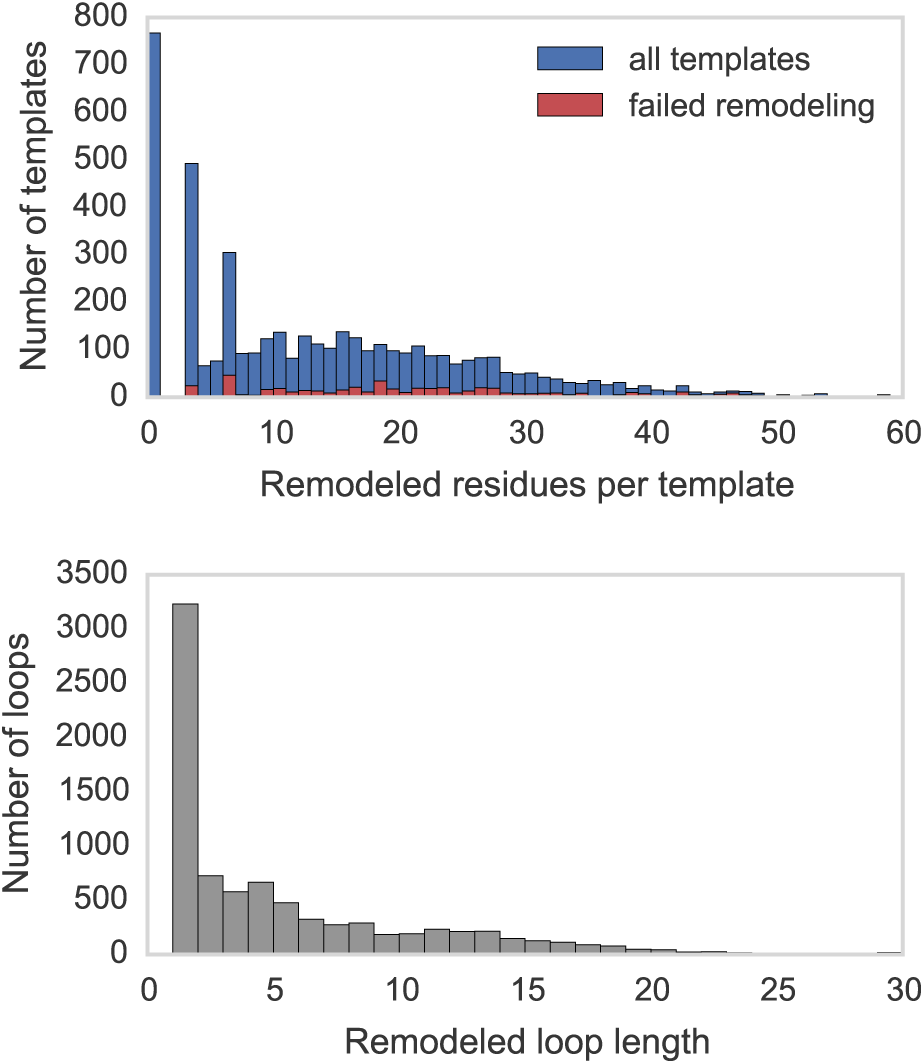
Distributions for the number of missing residues in the TK templates. The upper histograms show the number of missing residues per template, for all templates (blue) and for only those templates for which template remodeling with the loopmodel subcommand failed (red). The lower histogram shows the number of residues in each missing loop, for all templates.

Following loop remodeling, the **Ensembler** pipeline was performed up to and including the implicit solvent MD refinement stage, which completed with 389,067 (94%) surviving models across all TKs. To obtain statistics for the solvation stage without generating a sizeable amount of coordinate data (with solvated PDB coordinate files taking up about 0.9 MB each), the `solvate` subcommand was performed for two representative individual kinases *(Src* and *Abl1).*

The number of models which survived each stage are shown in Fig. 1, indicating that the greatest attrition occurred during the modeling stage. The number of refined models for each target ranged from 4046 to 4289, with a median of 4185, mean of 4184, and standard deviation of 57. Fig. 1 also indicates the typical timing achieved on a cluster for each stage, showing that the `build_models` and refine_implicit_md stages are by far the most compute intensive.

The files generated for each model (up to and including the implicit solvent MD refinement stage) totaled ∼116 kB in size, totalling 0.5 GB per TK target or42 GB for all 93 targets. The data generated per model breaks down as 39 kB for the output from the modelling stage (without saving MODELLER restraints files, which are about 397 kB per model) and 77 kB for the implicit solvent MD refinement stage.

### Evaluation of model quality and utility

#### All tyrosine kinases

To evaluate the variety of template sequence similarities relative to each target sequence, we calculated sequence identity distributions, as shown in Fig. 3. This suggests an intuitive division into three categories, with 355,712 models in the 0-35% sequence identity range, 51,330 models in the 35-55% range, and 5227 models in the 55-100% range. We then computed the RMSD distributions for the models created for each target (relative to the model derived from the template with highest sequence identity) Fig. 4, to assess the diversity of conformations captured by the modeling pipeline. Furthermore, to understand the influence of sequence identity on the conformational similarities of the resulting models, the RMSD distributions were stratified based on the three sequence identity categories described above. This analysis indicates that higher sequence identity templates result in models with lower RMSDs, while templates with remote sequence identities result in larger RMSDs on average.

**FIG. 3.**
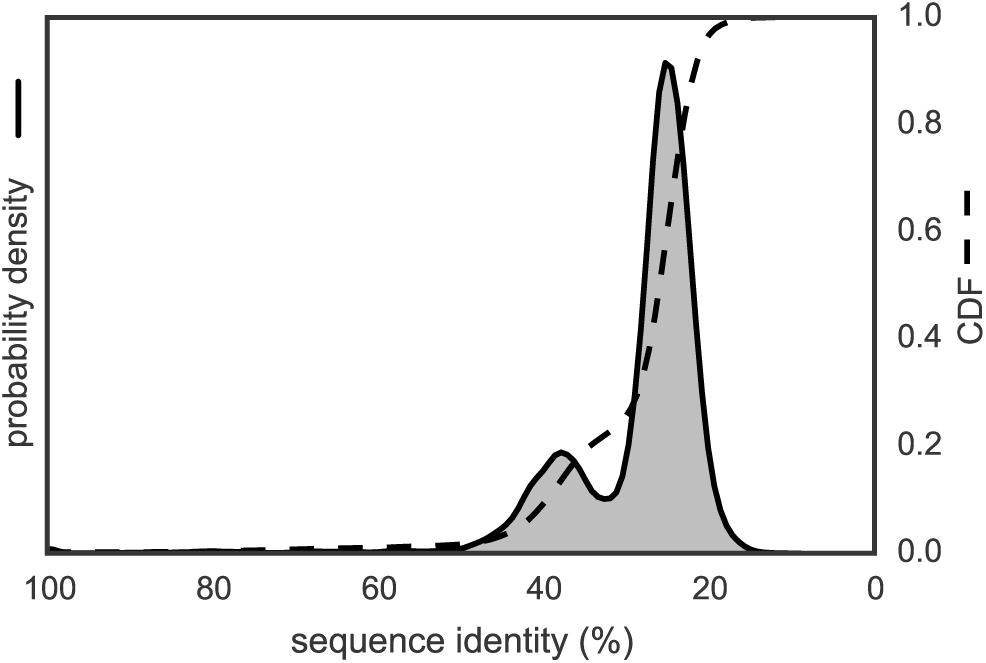
Template-target sequence identity distribution for human tyrosine kinase catalytic domains. Sequence identities are calculated from all pairwise target-template alignments, where targets are human kinase catalytic domain sequences and templates are all kinase catalytic domains from any organism with structures in the PDB, as described in the text. A kernel density estimate of the target-template sequence identity probability density function is shown as a solid line with shaded region, while the corresponding cumulative distribution function is shown as a dashed line.

**FIG. 4.**
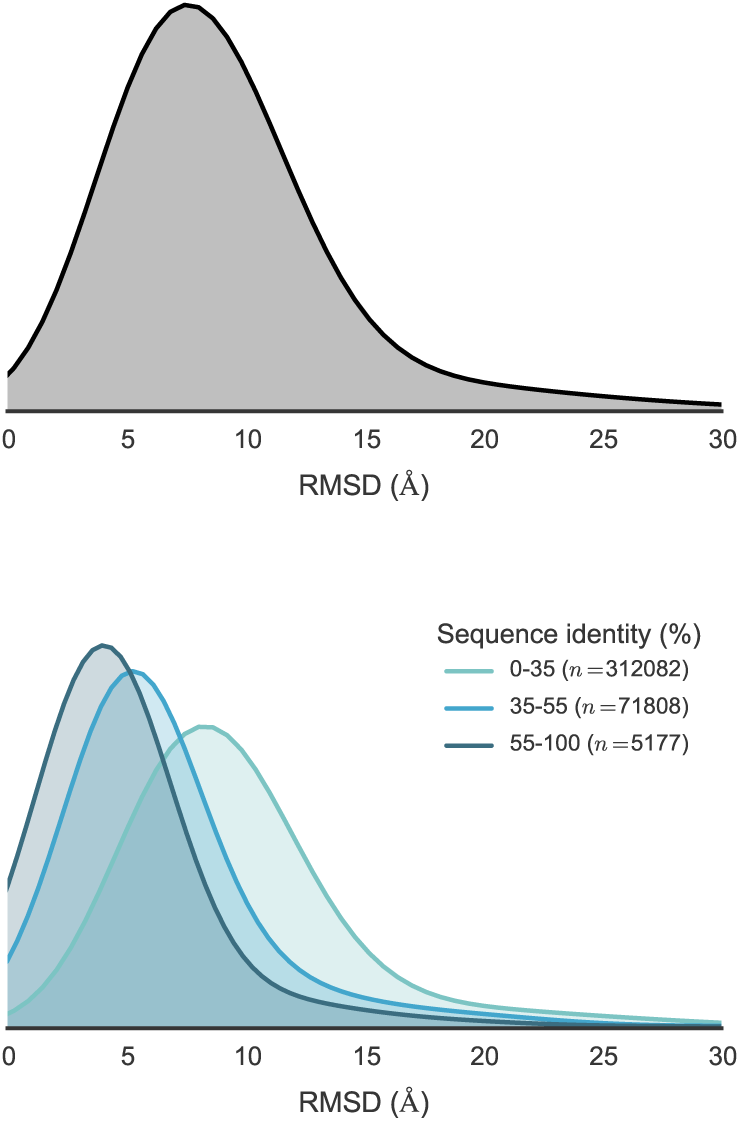
Distribution of RMSDs to allTK catalytic domain models relative to the model derived from the highest sequence identity template. Distributions are built from data from all 93 TK domain targets. To better illustrate how conformational similarity depends on sequence identity, the lower plot illustrates the distributions as stratified into three sequence identity classes: high identity (55-100%), moderate identity (35-55%), and remote identity (0-35%). The plotted distributions have been smoothed using kernel density estimation.

We also analyzed the potential energies of the models at the end of the implicit solvent MD refinement stage. These ranged from -14180 kT to -3160 kT, with a median of -9501 kT, mean of -9418 kT, and a standard deviation of 1198 kT (with a simulation temperature of 300 K). The distributions—stratified using the same sequence identity ranges as above—are plotted in Fig. 5, indicating that higher sequence identity templates tend to result in slightly lower energy models. Of the 4973 models which failed to complete the implicit refinement MD stage, all except 9 failed within the first 1 ps of simulation.

**FIG. 5.**
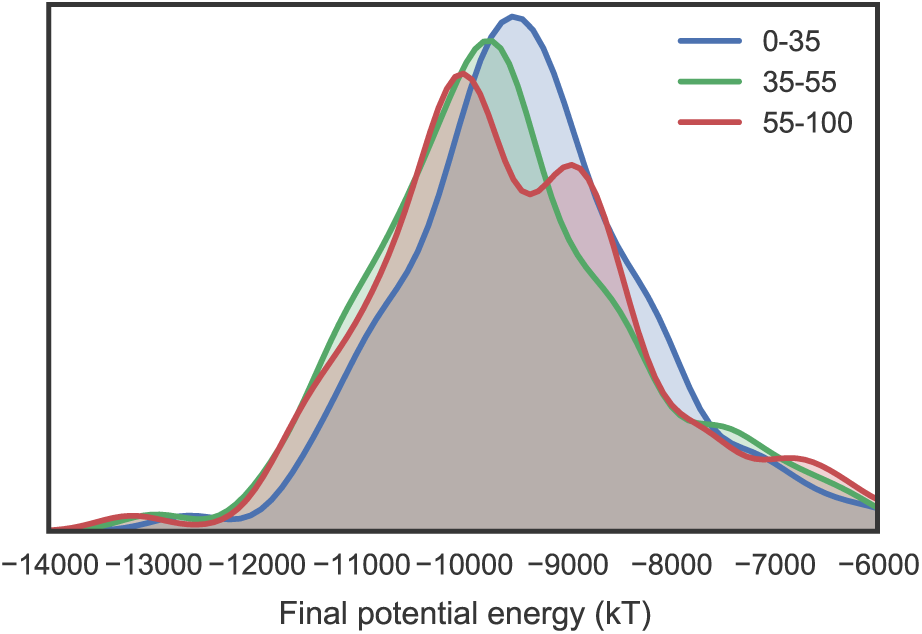
Distribution of final energies from implicit solvent MD refinement of TK catalytic domain models. To illustrate how the energies are affected by sequence identity, the models are separated into three sequence identity classes: high identity (55-100%), moderate identity (35-55%), and remote identity (0-35%). The plotted distributions have been smoothed using kernel density estimation. Refinement simulations were carried out at the default temperature of 300 K.

##### Src and Abl1

To provide a more complete evaluation of the models generated, we have analyzed two example TKs *(Src* and *Abl1)* in detail. Due to their importance in cancer, these kinases have been the subject of numerous studies, encompassing many different methodologies. In terms of structural data, a large number of crystal structures have been solved (with or without ligands such as nucleotide substrate or inhibitor drugs), showing the kinases in a number of different conformations. These two kinases are thus also interesting targets for MSM studies, with one recent study focusing on modeling the states which constitute the activation pathway of Src [45].

Fig. 6 shows a superposition of a set of representative models of Src and *Abl1.* Models were first stratified into three ranges, based on the structure of the sequence identity distribution (Fig. 3), then subjected to RMSD-based k-medoids clustering (using the msm builder clustering package [14]) to pick three representative models from each sequence identity range. Each model is colored and given a transparency based on the sequence identity between the target and template sequence. The figure gives an idea of the variance present in the generated models. High sequence identity models (in opaque blue) tend to be quite structurally similar, with some variation in loops or changes in domain orientation.

**FIG. 6.**
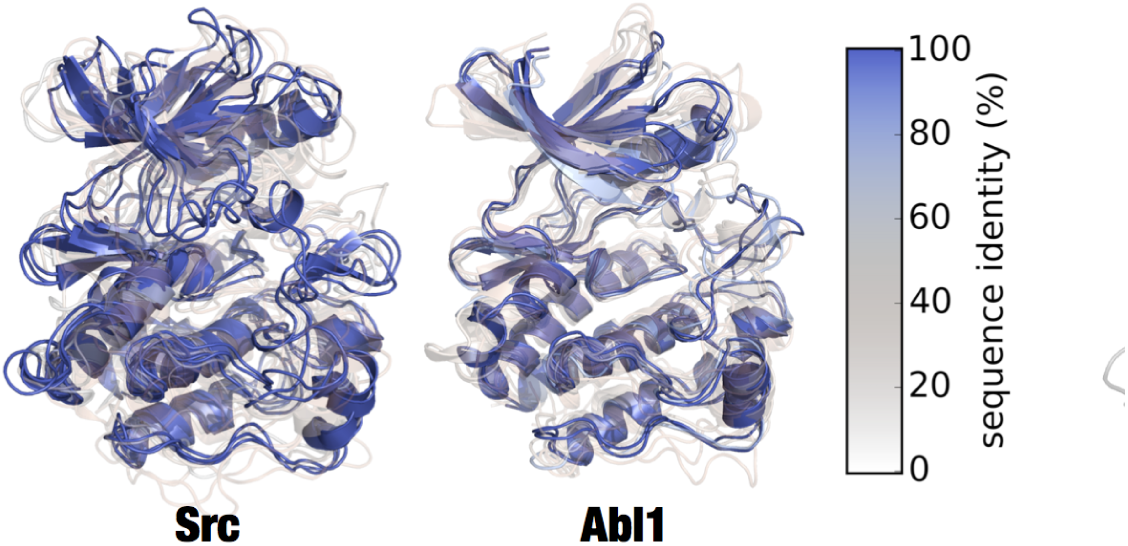
Superposition of clustered models of Src and Abl1. Superposed renderings of nine models each for Src and Abl1, giving some indication the diversity of conformations generated by Ensembler. The models for each target were divided into three sequence identity ranges (as in Fig. 4), and RMSD-based k-medoids clustering was performed (using the msmbuilder clustering package **[14]**) to select three clusters from each. The models shown are the centroids of each cluster. Models are colored and given transparency based on their sequence identity, so that high sequence identity models are blue and opaque, while lower sequence identity models are transparent and red.

The Abl1 renderings in Fig. 6 indicate one high sequence identity model with a long unstructured region at one of the termini, which was unresolved in the original template structure. While such models are not necessarily incorrect or undesirable, it is important to be aware of the effects they may have on production simulations performed under periodic boundary conditions, as long unstructured termini can be prone to interact with a protein’s periodic image. Lower sequence identity models (in transparent white or red) indicate much greater variation in all parts of the structure. We believe the mix of high and low sequence identity models to be particularly useful for methods such as MSM building, which require thorough sampling of the conformational landscape. The high sequence identity models could be considered to be the most likely to accurately represent true metastable states. Conversely, the lower sequence identity models could be expected to help push a simulation into regions of conformation space which might take intractably long to reach if starting a single metastable conformation.

To evaluate the models of Src and *Abl1* in the context of the published structural biology literature on functionally relevant conformations, we have focused on two residue pair distances thought to be important for the regulation of protein kinase domain activity. We use the residue numbering schemes for chicken Src (which is commonly used in the literature even in reference to human Src) [46, 47] and human Abl1 isoform A [48 -50] respectively; the exact numbering schemes are provided in Appendix 1.

Fig. 7 shows two structures of Src believed to represent inactive (PDB code: 2SRC) [46] and active (PDB code: 1Y57) [47] states. One notable feature which distinguishes the two structures is the transfer of an electrostatic interaction of E310 from R409 (in the inactive state) to K295 (in the active state), brought about by a rotation of the aC-helix. These three residues are also well conserved [51], and a number of experimental and simulation studies have suggested that this electrostatic switching process plays a role in a regulatory mechanism shared across the protein kinase family [45, 52, 53]. As such, we have projected the **Ensembler** models for Src and *Abl1* onto a space consisting of the distances between these two residue pairs (Fig. 8). The models show strong coverage of regions in which either of the electrostatic interactions is fully formed (for models across all levels of target-template sequence identity), as well as a wide range of regions in-between (mainly models with low sequence identity). We thus expect that such a set of models, if used as starting configurations for highly parallel MD simulation, could greatly aid in sampling of functionally relevant conformational states.

**FIG. 7.**
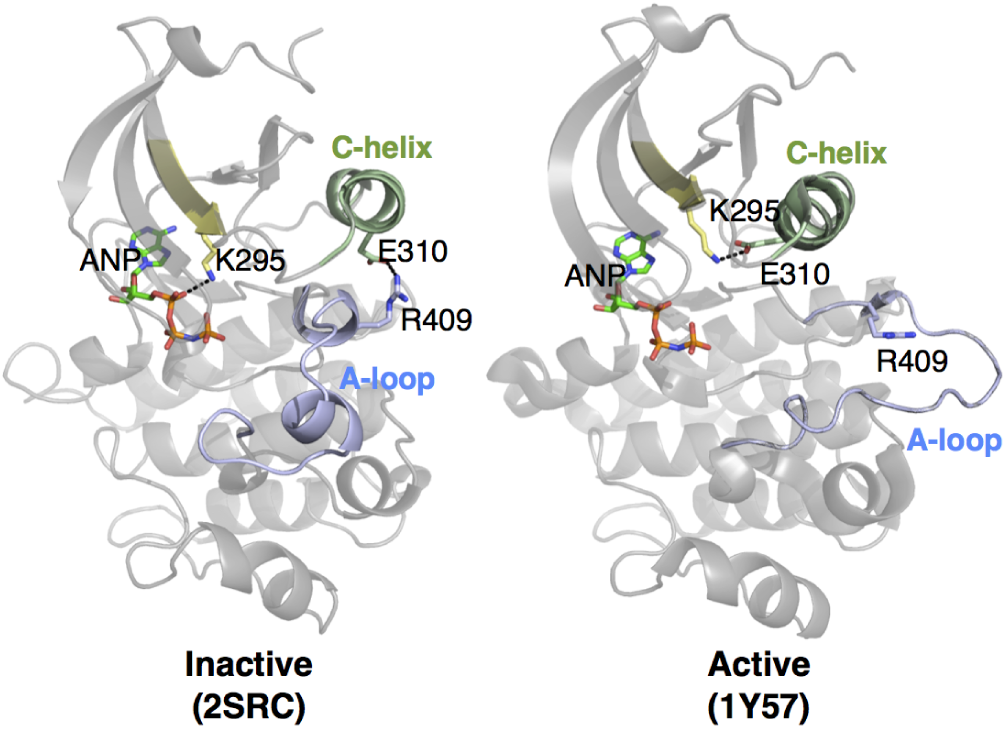
Two structures of Src, indicating certain residues involved in activation. In the inactive state, E310 forms a salt bridge with R409. During activation, the αC-helix (green) moves and rotates, orienting E310 towards the ATP-binding site and allowing it to instead form a salt bridge with K295. This positions K295 in the appropriate position for catalysis. Note that ANP (phospho-aminophosphonic acid-adenylate ester; an analog of ATP) is only physically present in the 2SRC structure. To aid visualization of the active site in 1Y57, it has been included in the rendering by structurally aligning the surrounding homologous protein residues.

**FIG. 8.**
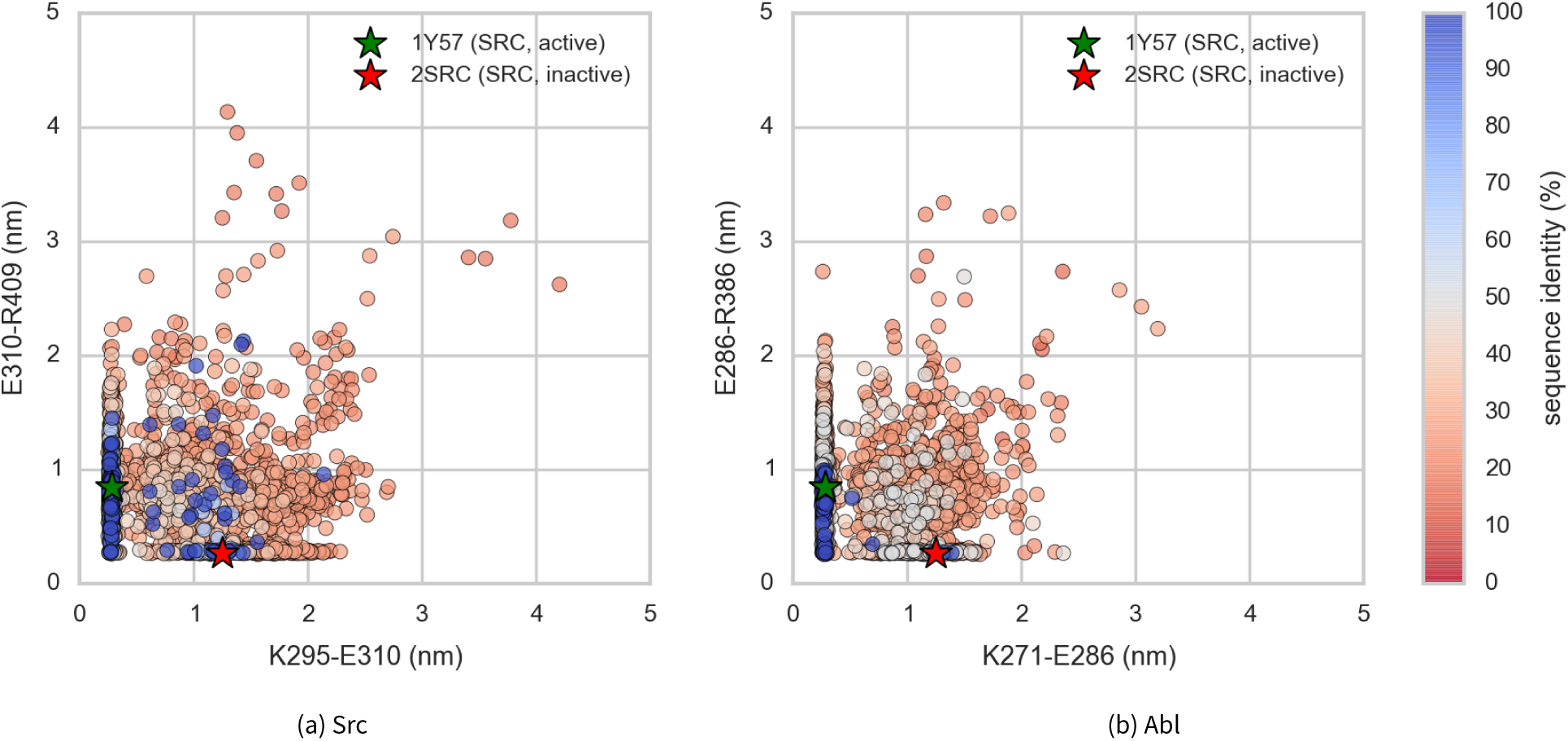
Src and Abl1 models projected onto the distances between two conserved residue pairs, colored by sequence identity. Two Src structures (PDB entries 1Y57 [47] and 2SRC [46]) are projected onto the plots for reference, representing active and inactive states respectively. These structures and the residue pairs analyzed here are depicted in Fig. 7. Distances are measured between the center of masses of the three terminal sidechain heavy atoms of each residue. The atom names for these atoms, according to the PDB coordinate files for both reference structures, are—Lys: NZ, CD, CE (ethylamine); Glu: OE1, CD, OE2 (carboxylate); Arg: NH1, CZ, NH2 (part of guanidine).

## IV. AVAILABILITY AND FUTURE DIRECTIONS

### Availability

The code for **Ensembler** is hosted on the collaborative open source software development platform GitHub (github.com/choderalab/ensembler). The latest release can be installed via the **conda** package manager for Python (conda.pydata.org), using the two commands shown in Box 2. This will install all dependencies except for MODELLER and Rosetta, which are not available through the **conda** package manager, and thus must be installed separately by the user. The latest source can be downloaded from the GitHub repository, which also contains up-to-date instructions for building and installing the code. Documentation can be found at ensembler.readthedocs.org.

#### Box 2.

##### Ensembler installation using conda.

~~~
conda config -add channels https://conda.binstar.org/omnia
conda install ensembler
~~~

A supplementary dataset can also be downloaded from the Dryad Digital Repository (DOI: 10.5061/dryad.7fg32). This contains the TK models described in the III section, general information on the targets and templates, plus a script and instructions for regenerating the same dataset.

### Future Directions

Comparative protein modeling and MD simulation set-up can be approached in a number of different ways, with varying degrees of complexity, and there are a number of obvious additions and improvements which we plan to implement in future versions of **Ensembler.**

Some amino acids can exist in different protonation states, depending on pH and on their local environment. These protonation states can have important effects on biological processes. For example, long time scale MD simulations have suggested that the conformation of the DFG motif of the TK Abl1—believed to be an important regulatory mechanism [54]—is controlled by protonation of the aspartate [55]. Currently, protonation states are assigned simply based on pH (a user-controllable parameter). At neutral pH, histidines have two protonation states which are approximately equally likely, and in this situation the selection is therefore made based on which state results in a better hydrogen bond. It would be highly desirable to instead use a method which assigns amino acid protonation states based on a rigorous assessment of the local environment. We thus plan to implement an interface and command-line function for assigning protonation states with MCCE2 [56-58], which uses electrostatics calculations combined with Monte Carlo sampling of side chain conformers to calculate pKa values.

Many proteins require the presence of various types of non-protein atoms and molecules for proper function, such as metal ions (e.g. Mg+^2^), cofactors (e.g. ATP) or posttranslational modifications (e.g. phosphorylation, methylation, glycosylation, etc.), and we thus plan for **Ensembler** to eventually have the capability to include such entities in the generated models. Binding sites for metal ions are frequently found in proteins, often playing a role in catalysis. For example, protein kinase domains contain two binding sites for divalent metal cations, and display significantly increased activity in the presence of Mg^2+^ [59], the divalent cation with highest concentration in mammalian cells. Metal ions are often not resolved in experimental structures of proteins, but by taking into account the full range of available structural data, it should be possible in many cases to include metal ions based on the structures of homologous proteins. We are careful to point out, however, that metal ion parameters in classical MD force fields have signif-icant limitations, particularly in their interactions with proteins [60]. Cofactors and post-translational modifications are also often not fully resolved in experimental structures, and endogenous cofactors are frequently substituted with other molecules to facilitate experimental structural analysis. Again, **Ensembler** could exploit structural data from a set of homologous proteins to model in these molecules, although there will likely be a number of challenges to overcome in the design and implementation of such functionality.

Another limitation with the present version of **Ensembler** involves the treatment of members of a protein family with especially long residue insertions or deletions. For example, the set of all human protein kinase domains listed in UniProt have a median length of 265 residues (mean 277) and a standard deviation of 45, yet the minimum and maximum lengths are 102 and 801 respectively. The latter value corresponds to the protein kinase domain of serine/threonine-kinase *greatwall,* which includes a long insertion between the two main lobes of the catalytic domain. In principle, such insertions could be excluded from the generated models, though a number of questions would arise as to how best to approach this.

## Conclusion

We believe **Ensembler** to be an important first step toward enabling computational modeling and simulation of proteins on the scale of entire protein families, and suggest that it could likely prove useful for tasks beyond its original aim of providing diverse starting configurations for MD simulations. The code is open source and has been developed with extensibility in mind, in order to facilitate its customization for a wide range of potential uses by the wider scientific community.

## V. ACKNOWLEDGMENTS

The authors are grateful to Robert McGibbon (Stanford) and Arien S. Rustenburg (MSKCC) for many excellent software engineering suggestions. The authors thank Nicholas M. Levinson (University of Minnesota), Markus A. Seeliger (Stony Brook), Diwakar Shukla (Stanford), and Avner Schlessinger (Mount Sinai) for helpful scientific feedback on modeling kinases. The authors are grateful to Benjamin Webb and AndrejSali (UCSF) for help with the MODELLER package, Peter Eastman and Vijay Pande (Stanford) for assistance with OpenMM, and Marilyn Gunner(CCNY) for assistance with MCCE2. All authors acknowledge support from the Sloan Kettering Institute. JDC, KAB, and DLP acknowledge partial support from NIH grant P30 CA008748. JDC and DLP also acknowledge the generous support of a Louis V. Gerstner Young Investigator Award. KAB was also supported in part by Starr Foundation grant I8-A8-058. PBG acknowledges partial funding support from the Weill Cornell Graduate School of Medical Sciences.

## Appendix 1: Sequences and residue numbering schemes for Src and Abl1

Kinase catalytic domains are highlighted in red, and the conserved residues analyzed in the main text (Figs. 7 and 8) are highlighted with yellow background.

### Human Abl1 sequence

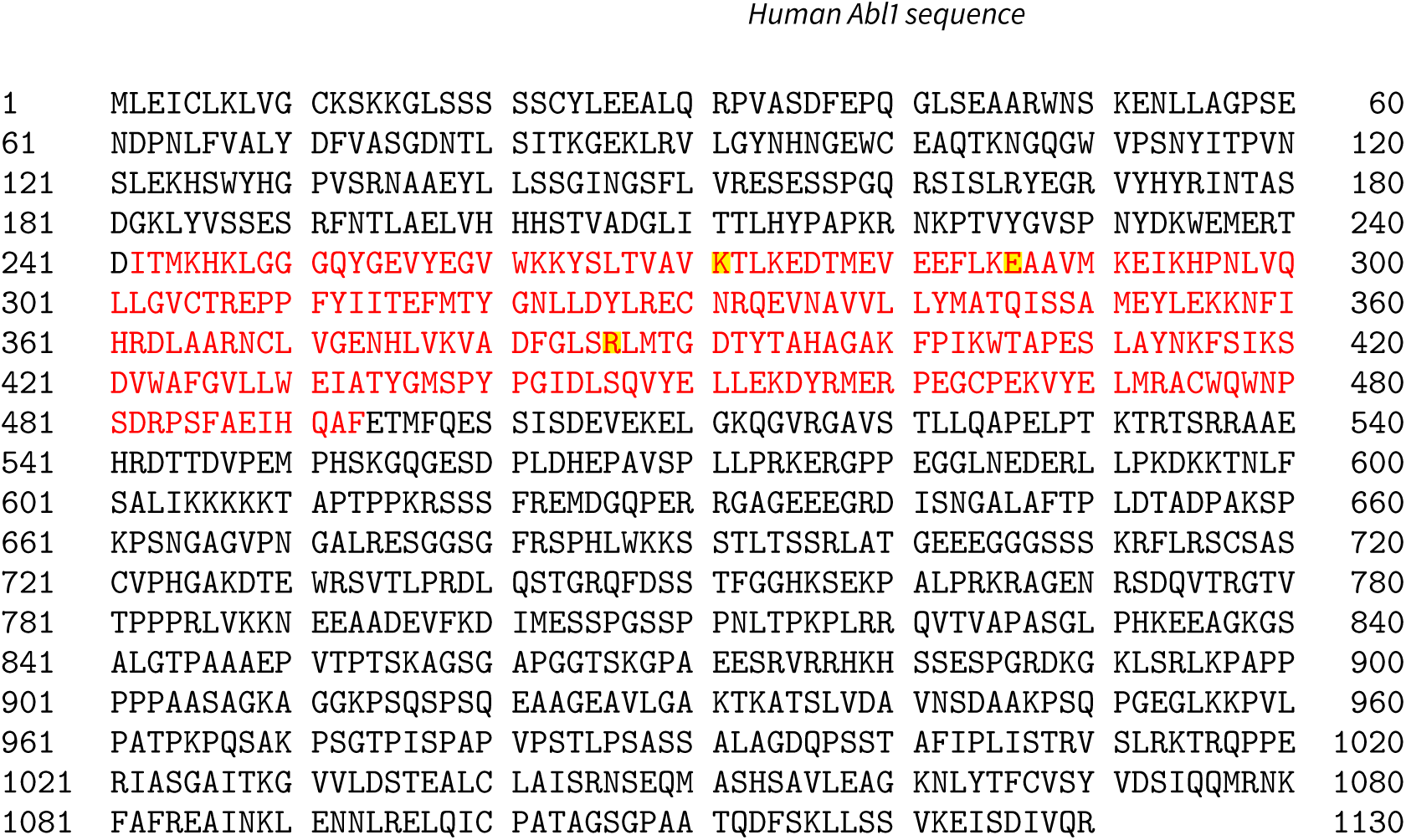

### Sequences for human and chicken Src, aligned using Clustal Omega

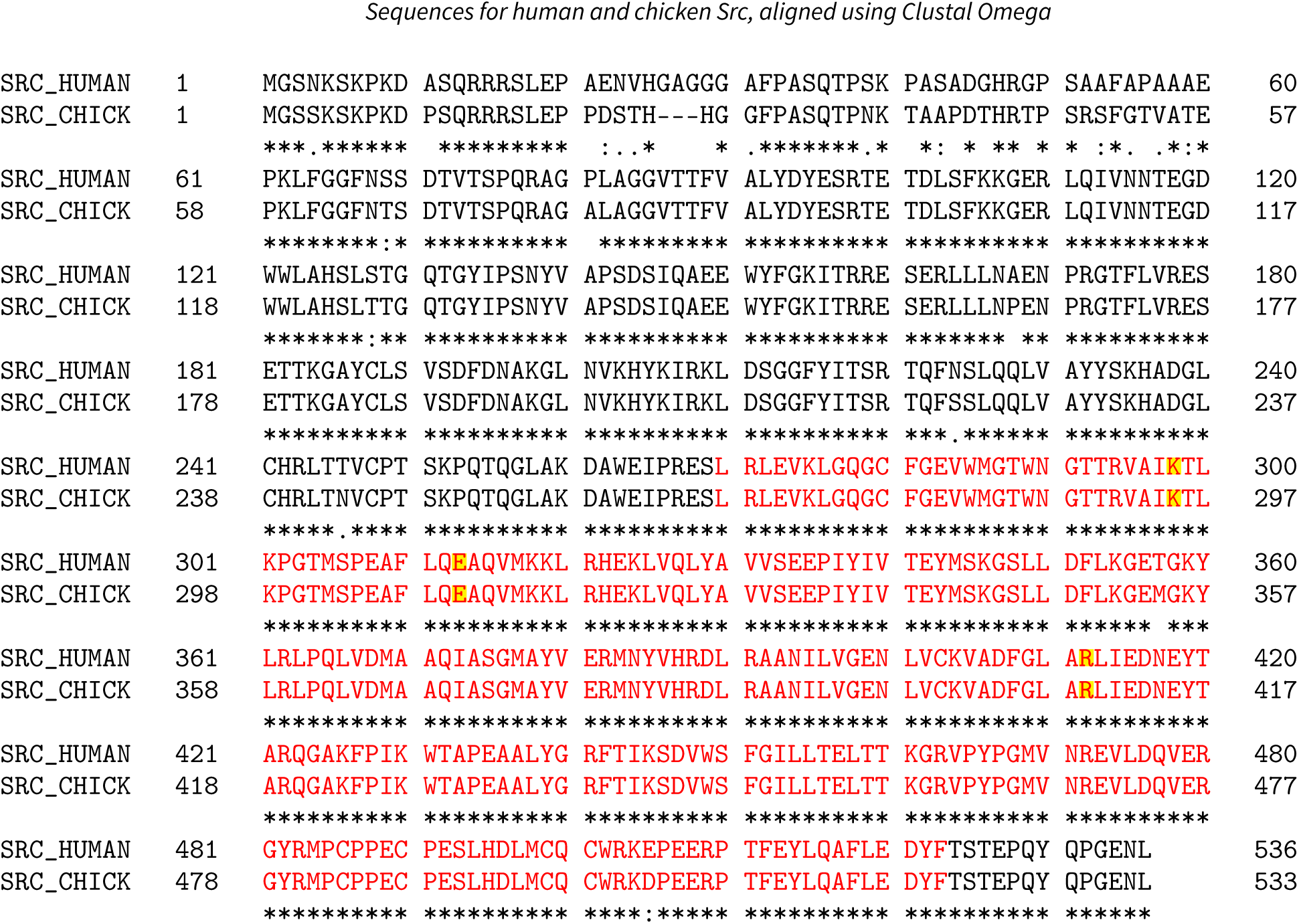

